# Punch in the gut: Parasite tolerance of phytochemicals reflects host diet

**DOI:** 10.1101/2021.10.07.463547

**Authors:** Evan C Palmer-Young, Ryan S Schwarz, Yanping Chen, Jay D Evans

**Affiliations:** USDA-ARS Bee Research Lab, Beltsville, MD, USA; Department of Biology, Fort Lewis College, Durango, CO, USA

**Keywords:** plant secondary metabolites, infectious disease spillover, One Health, *Apis mellifera*, *Lotmaria passim*, *Crithidia mellificae*, *Leishmania*

## Abstract

Gut parasites of plant-eating insects are exposed to antimicrobial phytochemicals that can reduce infection. Trypanosomatid gut parasites infect insects of diverse nutritional ecologies as well as mammals and plants, raising the question of how host diet-associated phytochemicals shape parasite evolution and host specificity. To test the hypothesis that phytochemical tolerance of trypanosomatids reflects the chemical ecology of their hosts, we compared related parasites from honey bees and mosquitoes—hosts that differ in phytochemical consumption—and contrasted our results with previous studies on phylogenetically related, human-parasitic *Leishmania*.

We identified one bacterial and ten plant-derived substances with known antileishmanial activity that also inhibited honey bee parasites associated with colony collapse. Bee parasites exhibited greater tolerance of chrysin—a flavonoid found in nectar, pollen, and plant resin-derived propolis. In contrast, mosquito parasites were more tolerant of cinnamic acid—a product of lignin decomposition present in woody debris-rich larval habitats. Parasites from both hosts tolerated many compounds that inhibit *Leishmania*, hinting at possible trade-offs between phytochemical tolerance and mammalian infection. Our results implicate the phytochemistry of host diets as a potential driver of insect-trypanosomatid associations, and identify compounds that could be incorporated into colony diets or floral landscapes to ameliorate infection in bees.

**ORIGINALITY AND SIGNIFICANCE:** Trypanosomatid parasites of insects pose threats to beneficial insects, crop plants, and human health. Elucidation of the environmental factors that determine host-parasite specificity and transmission of trypanosomatids is essential to predict the spread and spillover potential of trypanosomatids in wild and agricultural ecosystems. Our comparisons between trypanosomatids of bees and mosquitoes implicate the phytochemical composition of host diets as a potential determinant of insect-trypanosomatid associations. To our knowledge, this is the first comparative study of phytochemical tolerance in insect trypanosomatids. As the first characterization of phytochemical tolerance among honey bee-associated trypanosomatids, our results also identify compounds that could ameliorate infection of managed bees with parasites of concern to wild pollinators and associated with colony collapse.

## INTRODUCTION

Antimicrobial phytochemicals protect plants from biotic antagonists, including microbial infection, and can also have antiparasitic effects in the animals that consume these compounds (Huffman, 1997; Lozano, 1998; Hartmann, 2007; de Roode *et al*., 2013). In turn, parasites must tolerate host immunological and diet-related antimicrobial defenses—including phytochemicals—to establish successful infection (de Roode and Hunter, 2018). Because parasites that infect the gut are directly exposed to host-ingested substances, chemical tolerance of gut parasites is likely to reflect the dietary chemistry of their hosts, with consequences for insect host-parasite specificity that parallel the phytochemical-driven ecology of insects and plants (Raguso *et al*., 2014).

Insect gut-associated trypanosomatid protozoa infect diverse hosts with widely differing ecologies and patterns of phytochemical exposure (Frolov *et al*., 2021), making them a useful system for comparative analysis of chemical tolerance. Among the most well-studied are the *Leishmania* species. These sandfly-vectored parasites cause disease in >2 million humans each year, with 10% of the world’s population at risk, and have a greater health burden than any human parasite besides malaria (McGwire and Satoskar, 2014). These cases include >200,000 infections with visceral leishmaniasis, which, if untreated, results in >90% patient mortality (McGwire and Satoskar, 2014; Steverding, 2017). Due to their clinical significance, *Leishmania* spp. have been studied intensively in a search for affordable and effective treatments for human infections (De Rycker *et al*., 2013), including exhaustive testing of plant extracts and their components against both mammal-and insect-associated parasite life stages (Anthony *et al*., 2005; Le *et al*., 2018). These studies have suggested new treatments for trypanosomatid-associated infections of humans (Brindha *et al*., 2021) and beneficial insects (Richardson *et al*., 2015; Palmer-Young *et al*., 2016), and established patterns of phytochemical tolerance against which other trypanosomatids of differing ecologies can be compared.

Although most studies on trypanosomatid-inhibiting phytochemicals have been justifiably aimed at identification of candidate treatments for human infections, plant chemistry can also affect infection of insects and plants with both insect-specific and insect-vectored trypanosomatids and other protozoa. For example, different plant sugar sources affected malaria (*Plasmodium falciparum*) infection intensity and transmission potential in mosquitoes (Hien *et al*., 2016), and feeding of sand flies (*Phlebotomus papatasi*) on castor bean plants (*Ricinus communis*) led to the death of intestinal *Leishmania major* (Schlein and Jacobson, 1994). Among beneficial pollinating insects, sunflower (*Helianthus annuus*) pollen profoundly increased resistance to *Crithidia bombi* infection in bumble bees (Giacomini *et al*., 2018). More modest and variable reductions in infection occur in bees fed isolated compounds in nectar (Biller *et al*., 2015; Richardson *et al*., 2015; Thorburn *et al*., 2015; Palmer-Young, Hogeboom, *et al*., 2017), but plant communities were nevertheless important determinants of bee infection in experimental mesocosms (Adler *et al*., 2020). Plant secondary compounds can affect plant resistance to insect-vectored *Phytomonas* parasites of plants as well (Medina *et al*., 2015).

Circumstantial evidence for the role of plant chemistry in trypanosomatid evolution and host specificity can be found in the trypanosomatid genus *Phytomonas*. These insect-vectored parasites of plants have an expanded genomic repertoire of carbohydrate metabolism but lack the functional Krebs cycle and electron transport chain used by other trypanosomatids for aerobic respiration (Tielens and Van Hellemond, 1998). These traits are thought to reflect the diverse carbohydrates found in plants and provide protection against cyanide-based plant chemical defenses (Porcel *et al*., 2014; Jaskowska *et al*., 2015). In addition, *Phytomonas* are resistant to the plant lectins that agglutinate related *Crithidia* and *Herpetomonas* species (Petry *et al*., 1987). This suggests that host nutritional ecology could be an important driver of insect-trypanosomatid parasite interactions, with parasites exhibiting tolerance to the plant compounds encountered with the greatest frequency and intensity in the host gut.

Among the trypanosomatids’ many known insect hosts, bees occupy a unique nutritional and biochemical niche, characterized by nearly exclusive dependence on floral nectar and pollen that could pose phytochemical barriers to gut trypanosomatid infection. Both nectar and pollen contain diverse secondary metabolites that shape plant-pollinator ecology and plant-microbe ecology (Heil, 2011; Huang *et al*., 2012; Junker and Keller, 2015; Rivest and Forrest, 2020; Stevenson, 2020). Pollen in particular contains high phytochemical concentrations, which generally exceed those in nectar by several orders of magnitude (Palmer-Young, Farrell, *et al*., 2019). Flavonoids are one class of antimicrobial compounds (Cushnie and Lamb, 2005) that are ubiquitous in both nectar and pollen, with concentrations in pollen often exceeding 1% of total dry matter (Serra Bonvehi *et al*., 2001; Palmer-Young, Farrell, *et al*., 2019). Honey bees are further distinguished by collection of secondary metabolite-rich plant resins (propolis), which often contain over 20% flavonoids by mass (Bonvehí and Coll, 1994). This material is considered an integral component of the colony’s defense against parasites and pathogens (Simone-Finstrom and Spivak, 2010) and is a powerful inhibitor of *Leishmania* (Siheri *et al*., 2016; Alotaibi *et al*., 2021). The honey bee genome, which contains an expanded repertoire of flavonoid-metabolizing enzymes, reflects adaptations to tolerate chronic flavonoid exposure (Johnson *et al*., 2018), as does the flavonoid-metabolizing activity of the gut microbiome, which flourishes in the presence of pollen (Kešnerová *et al*., 2017). If the specialized ecology of the honey bee is reflected in its own phytochemical tolerance and that of its microbiota, it seems plausible that bee parasites are similarly adapted to nectar and pollen flavonoids—in contrast to the susceptibility to these compounds found in related *Leishmania* (Tasdemir *et al*., 2006)—and that these phytochemical barriers to infection could contribute to the limited diversity of trypanosomatids found to date in honey bees (Schwarz *et al*., 2015).

Two trypanosomatids are currently recognized in honey bees, each found on multiple continents and associated with individual and colony mortality. *Crithidia mellificae* was initially discovered during the investigation of colony losses in Australia that could not be explained by any then-recognized infectious disease (Langridge, 1966; Langridge and McGhee, 1967). This species can infect sweat, bumble, and mason bees (*Halictus, Bombus*, and *Osmia*) genera as well as honey bees (Strobl *et al*., 2019; Ngor *et al*., 2020), with elevated mortality in honey bees and *Osmia cornuta* (Strobl *et al*., 2019; Gómez-Moracho *et al*., 2020). *Lotmaria passim* was not distinguished from *C. mellificae* until 2014, but is now considered the dominant parasite of honey bees worldwide (Schwarz *et al*., 2015; Arismendi *et al*., 2016; Stevanovic *et al*., 2016; Xu *et al*., 2018; Williams *et al*., 2019). Prevalence has exceeded 80% in several national surveys (vanEngelsdorp *et al*., 2009; Cornman *et al*., 2012; Ravoet *et al*., 2013; Arismendi *et al*., 2016; Waters, 2018), with infection associated with colony collapse in the USA, Belgium, and Japan (Cornman *et al*., 2012; Morimoto *et al*., 2013; Ravoet *et al*., 2013). Experimentally infected bees exhibit increased mortality and diminished nutritional status (Buendía-Abad *et al*., 2020; Gómez-Moracho *et al*., 2020; Liu *et al*., 2020). Although the subacute effects of these parasites have yet to be fully elucidated, infection with the related *C. bombi* in bumble bees results in impaired colony establishment and growth (Brown *et al*., 2003), increased mortality during starvation and overwintering (Brown *et al*., 2000; Fauser *et al*., 2017), diminished reproduction (Brown *et al*., 2003), and pollination-relevant behavioral changes (Gegear *et al*., 2005, 2006; Otterstatter *et al*., 2005), as well facilitating parasite spillover to vulnerable sympatric bee species (Colla *et al*., 2006; Schmid-Hempel *et al*., 2014). *Crithidia bombi* showed high tolerance to many antileishmanial floral phytochemicals and even stimulation of growth by flavonoid-rich pollen extracts (Palmer-Young *et al*., 2016; Palmer-Young and Thursfield, 2017). However, the chemical sensitivity of the parasites in honey bees remains unknown. Comparing the phytochemical tolerance of trypanosomatids from bees with those of ecologically dissimilar hosts would provide insight into the role of plant secondary metabolites in structuring insect host-trypanosomatid parasite specificity, while also identifying compounds and forage plants with potential to manage infection in bee colonies.

Using the voluminous literature on *Leishmania*-inhibiting phytochemicals as a reference, we contrasted the phytochemical tolerance of these bee trypanosomatids with two strains of the type species of *Crithidia*—*C. fasciculata*. This *Crithidia* infects mosquitoes (Wallace, 1943) but is similar to bee trypanosomatids in morphology, intestinal niche, and fecal-oral transmission (Wallace, 1943; Clark *et al*., 1964). Although mosquitoes do consume sugar-based meals (including nectar) as adults (Hien *et al*., 2016), the type host of *C. fasciculata* (*Culex pipiens*) relies on blood—rather than pollen—as a nitrogen source. However, *C. fasciculata* infects larvae as well as adults (Wallace, 1943). As a result, the parasite is likely exposed to a variety of phytochemicals in the woody debris-rich aquatic breeding habitats of mosquitoes, which are considered to be the major sites of *C. fasciculata* transmission (Clark *et al*., 1964), and has been proposed as a chemically resistant model for trypanosomatid protozoa that infect mammals (Tasanor *et al*., 2006; Siheri *et al*., 2016), with which its phytochemical tolerance is correlated (Siheri *et al*., 2016).

Whereas positive selection for tolerance to nectar compounds is likely to predominate in parasites restricted to nectar-feeding insects, selection for secondary metabolite tolerance could be relaxed or even reversed in *Leishmania*, where the intracellular mammalian stage is shielded from direct exposure to plant metabolites. Previous authors have noted the lower tolerance of *Leishmania* and *Trypanosoma* to secondary metabolite-rich propolis relative to *C. fasciculata* (Siheri *et al*., 2016). Similarly, phylogenetic reconstructions have revealed an absence of catalase—an enzyme that mitigates the oxidative stress caused by flavonoid exposure (Fonseca-Silva *et al*., 2013)—in all trypanosomatids that infect mammals (Kraeva *et al*., 2017). This contrasts with the known or inferred presence of catalase in the insect-restricted species of *Crithidia, Lotmaria*, and their last common ancestor with *Leishmania* (Kraeva *et al*., 2017). If infection of mammals involved compromises in *Leishmania* tolerance to nectar secondary metabolites, these trypanosomatids could be more susceptible to the effects of plant compounds than their insect-restricted relatives.

To determine the potential for host phytochemical ecology to impose selective pressures on trypanosomatid parasites of insects, we compared tolerance of 25 known antileishmanial treatments— including flavonoids found at high concentrations in pollen and cinnamic acids associated with decay of woody debris—among phylogenetically related trypanosomatid parasites of honey bees (*C. mellificae* and *L. passim*) and mosquitoes (*C. fasciculata*). We predicted that parasites of honey bees—for whom the main source of proteins and lipids is floral pollen—would have higher tolerance of pollen-associated flavonoids than would parasites of mosquitoes, which obtain proteins and lipids from animal blood in the adult stage. In contrast, we predicted that parasites of mosquitoes would have greater tolerance of cinnamic acids associated with lignin breakdown in woody debris-rich larval habitats. Because all three parasite species use nectar-feeding insects as their sole hosts, we predicted that the tested parasites would be generally more tolerant to common nectar compounds than would the insect-vectored *Leishmania*, in which the intracellular mammalian stage is characterized by lesser phytochemical exposure. Building on prior research with *Leishmania*, our results identify eleven potential treatments for trypanosomatid infection in honey bee colonies, and implicate the phytochemical exposure of hosts as a potential driver of host-parasite specificity in insect trypanosomatids.

## MATERIALS AND METHODS

### Cell Cultures

*Crithidia mellificae* (ATCC 30254 (Langridge and McGhee, 1967)), *L. passim* (strain BRL (Schwarz *et al*., 2015)) and *C. fasciculata* strains “CFC1” (from Michael & Megan Povelones (Filosa *et al*., 2019)) and “Wallace” (ATCC 12857) were obtained from the American Type Culture Collection and collaborators. Honey bee parasites were grown in ‘FPFB’ medium including 10% heat-inactivated fetal bovine serum (pH 5.9-6.0) (Salathé *et al*., 2012). Mosquito parasites were grown in brain-heart infusion broth with 20 ug/mL hemin (pH 7.4). All parasites were incubated at 20 °C in vented cell culture flasks and transferred to fresh media every 2 d (Palmer-Young *et al*., 2021).

### Chemical treatments

To identify parasite-inhibiting compounds and compare patterns of tolerance between trypanosomatids of honey bees and mosquitoes, we tested the concentration-dependent effects of 25 treatments on growth rates of parasite cell cultures. The compounds were chosen based on previously demonstrated inhibitory activity against *Leishmania* and, in the case of methyl jasmonate, related parasites (Gold *et al*., 2003; Ofer *et al*., 2008). We tested seven flavonoids (five aglycones and 2 glycosides), five phenylpropanoids (including three cinnamic acids), three benzenoids, three terpenoids, two alkaloids, one oxylipin, and three plant extracts. As a positive control for inhibition, we used the polyene antifungal and antileishmanial drug amphotericin B **(Fig. 1, Supplementary Table 1)**.

**Figure 1.**
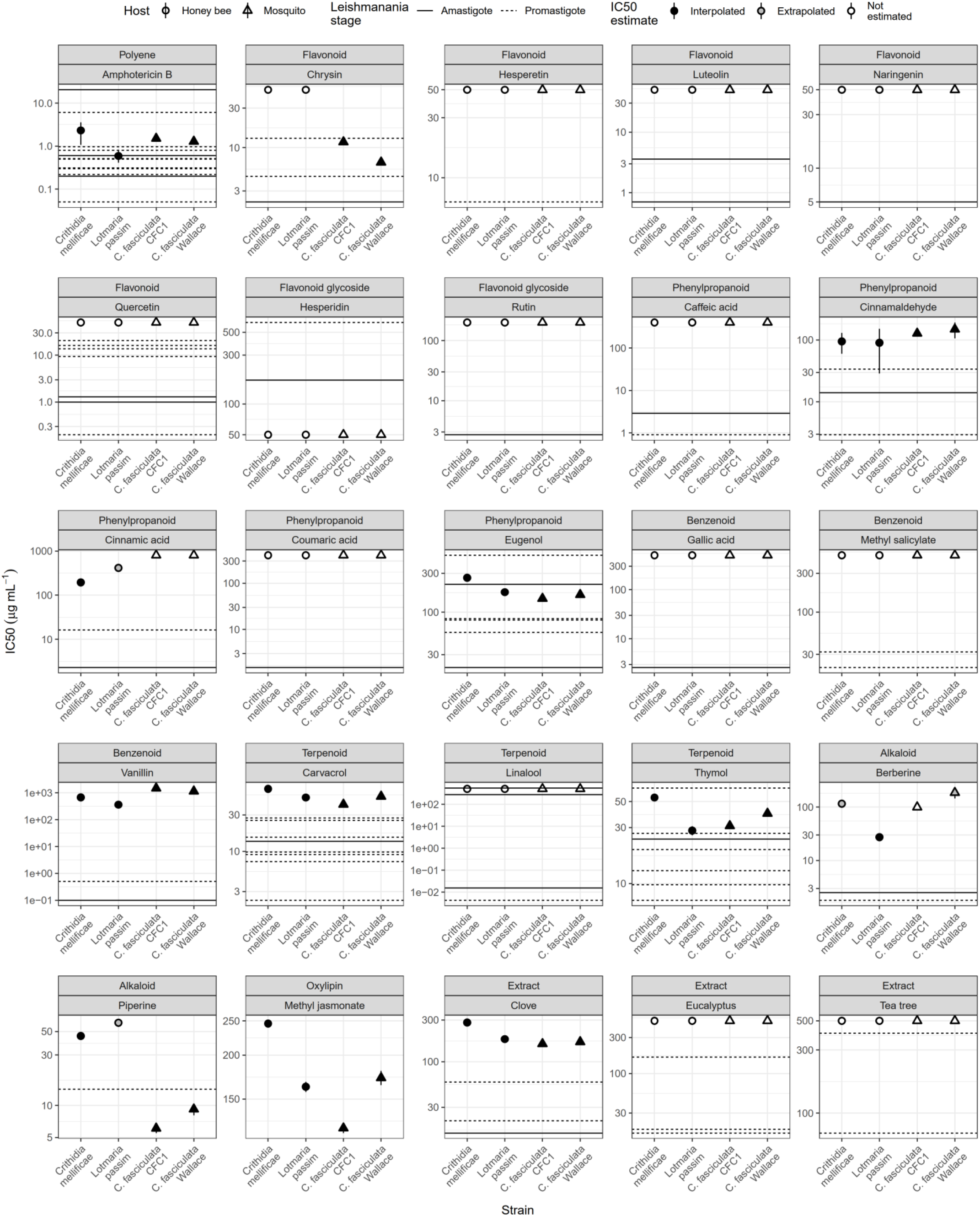
Chemical tolerance of trypanosomatid parasites from honey bees (*Crithidia mellificae* and *Lotmaria passim*, circles) and mosquitoes (*Crithidia fasciculata*, triangles), grouped by treatment type (top line of facet label). Y-axis show 50% Inhibitory treatment concentrations (IC50) for parasite growth rate. Points and error bars show estimates and 95% confidence intervals for the IC50. Shading of points indicates whether estimated IC50 was within the tested concentration range (“Interpolated”, black fill), above the tested range (“Extrapolated”, gray circles), or not estimated due to insufficient inhibition (unshaded points). In the latter case, points indicate the maximum concentration tested and no error bars are shown. Some error bars on estimated values are not visible because they are smaller than the corresponding points. Horizontal reference lines indicate published IC50 values for *Leishmania* spp. (solid lines: amastigote (i.e., mammalian) stage; dashed lines: promastigote (insect) stage). Dose-response curves used for IC50 estimation are shown in **Supplementary Figure 1**. Literature references for *Leishmania* IC50 estimates are given in **Supplementary Table 1**. Numerical IC50 estimates for the parasites tested here are given in **Supplementary Table 2**.

Compounds were dissolved in ethanol or DMSO (depending on solubility of compounds) prior to preparation of serial two-fold dilutions in growth media. Solvent concentrations in the final assay did not exceed 1% ethanol or 2% DMSO, each of which had minimal effects on parasite growth. Within each experiment, all treatment concentrations contained the same concentration of solvent.

### Phytochemical exposure experiments

Parasite growth rates were measured by optical density (OD_600_) on a microplate reader with 32°C incubation temperature. Cultures were diluted in fresh media and combined with equal volumes of 2x chemical treatments in 96-well plates for a starting OD of 0.020 in a volume of 100 µL media per well. Measurements were taken every 15 min for 48 h, with a 30 s shake before each read. A separate experiment was conducted for each parasite type (bee- or mosquito-associated). Each experiment included five replicates per strain at each of eight treatment concentrations (including a 0 µg ml^−1^control treatment), plus two cell-free control wells at each concentration to control for any growth-independent changes in OD during incubation. We selected the twelve compounds where the highest treatment concentration resulted in at least 50% reduction in OD after 24 h for dose-response modeling.

### Phytochemical tolerance analysis

Analyses were conducted using R for Windows v4.0.3 (R Core Team, 2014). Models were optimized using package “drc”(Ritz *et al*., 2015). Figures were made with packages “ggplot2” and “cowplot” (Wickham, 2009; Wilke, 2016).

#### Growth rates

Net OD was calculated by subtracting the average OD of cell-free controls of the corresponding media, treatment concentration, and time point. Maximum growth rate for each well within the first 24 h of incubation was computed based on the maximum slope of a rolling regression of ln(OD) vs. time, using a time window corresponding to approximately two cell divisions (5 h for *C. mellificae*, 12 h for *L. passim*, 10 h for *C. fasciculata* CFC1, 7 h for *C. fasciculata* Wallace) (Padfield *et al*., 2020). The first 2 h of each run were excluded to allow OD readings to stabilize. We used only slopes with r^2^ values of >0.80. Samples where mean growth rate was negative or where linear model r-squared values were <0.8 for all time windows (indicating growth indistinguishable from background noise) were assigned a maximum growth rate of zero.

#### Dose-response models

The relationship between growth rate and treatment concentration was modeled using a logistic regression with a lower limit of zero to estimate concentrations that inhibited growth by 50% (IC50) (Ritz *et al*., 2015; Palmer-Young, Raffel, *et al*., 2019). Strains were considered significantly different when the 95% confidence intervals for their IC50’s did not overlap. We used units of µg mL^−1^ to facilitate comparisons between pure compounds and extracts of undefined molarity.

## RESULTS

### Phytochemical tolerance of bee and mosquito parasites

#### Most inhibitory treatments

Although the tested substances were chosen based on demonstrated inhibition of *Leishmania* in previous work **(Supplementary Table 1)**, inhibition was sufficient to estimate dose-response models for only 11 treatments each for the parasites of honey bees and mosquitoes. Except for chrysin (no effect on bee parasites up to 50 μg mL^−1^) and cinnamic acid (minimal effect on mosquito parasites up to 800 μg mL^−1^), the same treatments inhibited growth of both parasite pairs. For the remaining treatments, inhibitory concentrations were on the same order of magnitude across the two groups.

The three compounds with the strongest inhibitory effects (IC50 <100 μg mL^−1^ for all strains) were the antifungal and first-line antileishmanial *Streptomyces* metabolite amphotericin B (IC50 range 0.59 ± 0.089 μg mL^−1^ SE for *L. passim* to 2.32 ± 0.62 μg mL^−1^ for *C. mellificae*) and the terpenoids thymol (28.3 ± 1.22 μg mL^−1^ for *L. passim* to 54.1 ± 0.84 μg mL^−1^ for *C. mellificae*) and its isomer carvacrol (41.1 ± 1.91 μg mL^−1^ for *C. fasciculata* CFC1 to 65.7 ± 0.81 μg mL^−1^ for *C. mellificae*).

Dose-response curves were also estimable for the following compounds, in order of increasing minimum IC50: cinnamaldehyde (90.3 ± 30.4 μg mL^−1^ for *L. passim* to 149 ± 21.1 μg mL^−1^ for *C. fasciculata* Wallace), methyl jasmonate (124 ± 1.90 μg mL^−1^ for *C. fasciculata* CFC1 to 246 ± 2.21 μg mL^−1^ for *C. mellificae*), eugenol (147 ± 2.97 μg mL^−1^ for *C. fasciculata* CFC1 to 264 ± 2.65 μg mL^−1^ for *C. mellificae*), clove oil (160 ± 2.59 μg mL^−1^ for *C. fasciculata* CFC1 to 280 ± 6.82 μg mL^−1^ for *C. mellificae*), and vanillin (362 ± 11.3 μg mL^−1^ for *L. passim* to 1487 ± 12.0 μg mL^−1^ for *C. fasciculata* CFC1). The alkaloids berberine and piperine both showed inhibitory effects, but IC50 estimation was limited by compound solubility. Piperine had a stronger inhibitory effect (IC50 range 6.09 ± 0.29 μg mL^−1^ for *C. fasciculata* CFC1 to 60.6 ± 1.26 μg mL^−1^ for *L. passim*) than did berberine (27.4 ± 1.17 μg mL^−1^ for *L. passim* to 186 ± 20.3 μg mL^−1^ for *C. fasciculata* Wallace). Note that this last estimate is an approximation, as <30% inhibition was found at the highest concentration tested (100 μg mL^−1^) **(Supplementary Figure 1)**.

#### Differences between parasites from the same host

Among the two honey bee parasites, *C. mellificae* was more tolerant than *L. passim* of 8 of 11 inhibitory compounds, whereas *L. passim* showed greater tolerance of cinnamic acid and piperine **(Figure 1)**. For cinnamaldehyde, IC50’s of the two species were indistinguishable **(Figure 1)**. Differences between *C. mellificae* and *L. passim* exceeded 2-fold for cinnamic acid (414 ± 12.0 μg mL^−1^ for *L. passim* vs. 195 ± 5.20 μg mL^−1^ for *C. mellificae*) and 4-fold for berberine (116 ± 7.25 μg mL^−1^ for *C. mellificae* vs. 27.4 ± 1.17 μg mL^−1^ for *L. passim*). IC50 values for the two strains of *C. fasciculata* did not differ by >2-fold for any of the treatments **(Figure 1)**.

#### Differences between bee and mosquito parasites

Qualitative differences between honey bee and mosquito parasites were found for the flavonoid chrysin (IC50 >50 μg mL^−1^ for both *C. mellificae* and *L. passim* vs. 6.65 ± 0.23 μg mL^−1^ for *C. fasciculata* CFC1 and 11.7 ± 0.42 μg mL^−1^ for *C. fasciculata* Wallace), indicating >4-fold higher tolerance among the bee parasites. Bee parasites were also more tolerant of the alkaloid piperine (IC50 range 45.3-60.6 μg mL^−1^) than were the mosquito parasites (6.09-9.22 μg mL^−1^) **(Figure 1)**. In contrast, tolerance of cinnamic acid was >2-fold higher among the mosquito parasites (>800 μg mL^−1^, extrapolated IC50 range 1252-1637 μg mL^−1^ for *C. fasciculata* **(Supplementary Figure 1)**, vs. 195 ± 5.20 μg mL^−1^ for *C. mellificae* and 414 ± 12.0 μg mL^−1^ for *L. passim*). Tolerance of vanillin was likewise ∼2-fold higher among the mosquito parasites (1146-1487 μg mL^−1^) than the bee parasites (362-677 μg mL^−1^). For the other compounds, boundaries of the IC50 ranges differed by <2-fold between the two groups **(Figure 1)**.

#### High flavonoid tolerance relative to previously tested *Leishmania*

We found minimal inhibition by the remaining 13 treatments **(Figure 1)**. In contrast to their inhibitory effects against *Leishmania* (except for hesperidin), there was a conspicuous absence of inhibition by flavonoids and flavonoid glycosides. None of the seven tested glycosides or aglycones inhibited either parasite of honey bees, and only one inhibited the mosquito-associated *C. fasciculata*. A similar pattern was seen for the cinnamic acid-based phenylpropanoids (cinnamic, coumaric, and caffeic acids). The honey bee parasites were affected only by cinnamic acid, with the mosquito parasites unaffected by all three **(Figure 1)**. For the treatments where we found 6 or more literature references **(Supplementary Table 1)**, IC50 values of the parasites tested here were generally two-to four-fold higher relative to previous results with *Leishmania* (e.g., Amphotericin B: 2.8-fold (median 1.39 μg mL^−1^ for our parasites vs. 0.505 μg mL^−1^ for *Leishmania*, n = 12 references); carvacrol: 4.4-fold (51.5 vs. 11.7 μg mL^−1^, n = 10 references); thymol: 2.2-fold (35.3 vs 16.2 μg mL^−1^, n = 8 references); eugenol: 2.1-fold (170 vs 81.4 μg mL^−1^, n = 6 references). This indicates that the >10-fold elevated flavonoid tolerance of our parasites relative to *Leishmania* is not merely an artefact of methodological differences.

## DISCUSSION

Our study provides the first characterization of the direct effects of plant secondary metabolites on trypanosomatid parasites of honey bees, identifying potential treatments for globally prevalent infections that have been associated with colony loss. Comparisons with the mosquito parasite *C. fasciculata* demonstrate the broad potential for host diet-associated phytochemicals to shape insect host-trypanosomatid ecology; to our knowledge, this is the first comparative study of phytochemical tolerance in insect trypanosomatids.

### Parasite-inhibiting compounds and their applications

The most potent growth inhibitor of both bee and mosquito parasites was the antifungal and antileishmanial drug amphotericin B (Yardley and Croft, 1997), which targets cell membranes containing ergosterol—the main membrane sterol in both fungi and trypanosomatids (Xu *et al*., 2014). However, this compound also damages the membranes of insect cells (Millam Stanley and Vaughn, 1967). Because insects cannot synthesize sterols *de novo* to repair this damage, their cells are 20-to 40-fold more susceptible to amphotericin B than are mammal cells *in vitro* (Millam Stanley and Vaughn, 1967). Mosquito and moth cell lines exhibited pathological changes at concentrations of 1-2 μg mL^−1^— within the IC50 range of our parasites (0.59-2.32 μg mL^−1^)—suggesting that despite its potency against parasites, this compound might not be sufficiently selective for use in bees.

The inhibitory effects of thymol, carvacrol, and eugenol—as well as eugenol-dominated clove extract (Maggi *et al*., 2010)— are consistent with their well-documented activity against *Leishmania* ((Le *et al*., 2018), **Figure 1, Supplementary Table 1**). Our inhibitory concentrations are comparable to those obtained in previous work on other strains of *C. fasciculata* and the bumble bee parasite *C. bombi*. For thymol, the 28-54 μg mL^−1^ IC50 range for *L. passim* and *C. mellificae* is similar to the 8.5-49.8 μg mL^−1^ for four strains of *C. bombi* (Evan C. Palmer-Young, Sadd, *et al*., 2017), while the 31-40 μg mL^−1^ IC50 for *C. fasciculata* matches the 32.5 μg mL^−1^ reported previously (Azeredo and Soares, 2013). For eugenol, the 176-264 μg mL^−1^ IC50 range of the honey bee parasites overlaps with the 44-185 μg mL^−1^ IC50 range for *C. bombi* (Evan C. Palmer-Young, Sadd, *et al*., 2017), while the 147-164 μg mL^−1^ range for *C. fasciculata* is within two-fold of the previously calculated 93.7 μg mL^−1^ (Azeredo and Soares, 2013). The activity of these compounds against diverse trypanosomatids at similar concentrations suggests that they have conserved or, more likely, multiple cellular targets that ensure effectiveness against parasites, although slight increases in tolerance were seen over 6 weeks of chronic exposure in *C. bombi* (Evan C. Palmer-Young, Sadd, *et al*., 2017). The observation that both thymol and eugenol—like amphotericin B—appear to interfere with trypanosomatid membrane function (Santoro *et al*., 2007; Azeredo and Soares, 2013) hints that membranes might make good treatment targets for trypanosomatids of both insects and mammals.

If these inhibitory compounds prove effective against infection and non-toxic for bees, they could be incorporated into bee diets—either by providing forage plants with phytochemical-rich nectar and pollen, or deliberate supplementation of honey bee colonies—to improve resistance to trypanosomatids. For example, *T. vulgaris* nectar contains thymol (26.1 μg mL^−1^) at concentrations comparable to the IC50 for *Lotmaria* (28.3 μg mL^−1^). Regarding therapeutic hive treatments, comparisons with previous studies suggest that honey bees can tolerate both thymol and eugenol-rich clove oil at concentrations well above those needed to inhibit growth of gut parasites. For thymol, the 8 d LD50 (>1,000 μg mL^−1^ (Ebert *et al*., 2007)) is well above the 28-54 μg mL^−1^ IC50 range for the parasites *L. passim* and *C. mellificae*, implying a >20-fold margin of safety for medication of bees with this compound. For clove oil, the 8 d LD50 (7800 μg mL^−1^ (Ebert *et al*., 2007)) is similarly nearly 30-fold higher than the 181-280 μg mL^−1^ IC50 range for the honey bee trypanosomatids. However, these lipophilic compounds are rapidly absorbed from the intestine, with a half-life of ∼2 h and >90% absorbance with 8 h in pigs (Michiels *et al*., 2008). The extended gut transit time of newly emerged honey bees (12-24 h for ingesta to reach the hindgut, where parasites establish (Crailsheim, 1988; Gisder *et al*., 2020)) could effectively shield parasites from the effects of ingested phytochemicals (Palmer-Young *et al*., 2018), such that these compounds are less selective than the ratio between bee LD50 and parasite IC50 suggests. On the other hand, midgut retention time may be as little as 1 h for older male bees (Ferreira and Landim, 2003), implying that the hindgut concentrations and antiparasitic effects of ingested phytochemicals could vary by sex and increase with host age.

### Host ecology-associated differences between parasites of bees and mosquitoes

The high tolerance of the flavonoid chrysin in bee relative to mosquito parasites is consistent with the central role of flavonoids in bee ecology, evolution, and microbiome metabolism (Kešnerová *et al*., 2017; Johnson *et al*., 2018). Flavonoid concentrations in honey (42-212 μg g^−1^, with acacia honey having 0.47 μg g^−1^ chrysin) have been described as “copious” (Pichichero *et al*., 2009), but are dwarfed by the ∼5,000 μg g^−1^ in bee pollen (Serra Bonvehi *et al*., 2001). Flavonoid concentrations in propolis—to which parasites are likely exposed in the colony environment and on mouthparts—are 40-fold higher still (>20% by mass, or 200,000 μg g^−1^), including up to 4900 μg g^−1^ chrysin (i.e., four orders of magnitude higher than in the acacia honey (Bonvehí and Coll, 1994)). In contrast to the rapidly metabolized terpenoids and flavonoids, flavonoids are poorly absorbed (Hostetler *et al*., 2017) and readily detected in the honey bee intestinal lumen (Kešnerová *et al*., 2017). This ability of flavonoids to reach the hindgut intact could allow these compounds to exert selective pressure on hindgut-dwelling parasites such as *Crithidia* and *Lotmaria*. Although the mode of action of these compounds was not investigated here, one target of flavonoids in *Leishmania* is the enzyme arginase (da Silva *et al*., 2012). The genome of *L. passim* (originally published as *C. mellificae*) was noted for containing an arginase of putative bacterial origin (Runckel *et al*., 2014), whereas plant-parasitic *Phytomonas* lacked arginase activity entirely (Camargo, 1999), suggesting that further investigation into the role of this gene in tolerance of flavonoid-rich host environments is warranted.

The high tolerance of bee parasites to the alkaloid berberine is intriguing both ecologically and clinically. The effects on alkaloids on plant-pollinator interactions—including plant and pollinator reproduction (Kessler *et al*., 2012; Arnold *et al*., 2014) and coevolution (Adler *et al*., 2012) and bee physiology (du Rand *et al*., 2015), behavior (Wright *et al*., 2013), and infection (Manson *et al*., 2010; Köhler *et al*., 2012)—have received considerable attention. Alkaloids were the most abundant class of secondary metabolites in pollens—occurring at concentrations slightly exceeding those of flavonoids (Palmer-Young, Farrell, *et al*., 2019). Although testing of additional compounds is needed, our results suggest that the alkaloids may also influence the ecology and evolution of gut parasites in pollinivorous bees. Tolerance of piperine specifically is notable given recent interest in development of *Piper* spp. extracts and piperine-based treatments for *Leishmania* (Ferreira *et al*., 2011; Le *et al*., 2018). The honey bee trypanosomatid *L. passim* could serve as a piperine-resistant model to understand parasite adaptations that might affect the efficacy of such treatments.

The greater tolerance of the mosquito parasite *C. fasciculata* to two compounds associated with wood decomposition—cinnamic acid and vanillin (Higuchi, 1971)—suggests parasite adaptation to the aquatic habitats and food sources occupied and consumed by larvae, which is considered the most important stage for this parasite’s transmission (Clark *et al*., 1964). Mosquito larvae filter-feed on bacteria in the water column and browse directly on decaying leaves, wood, and other plant material (Merritt *et al*., 1992), in which total phenolic concentrations can reach 10,000-20,000 μg g^−1^ (Rey *et al*., 2000). Leachates from decaying leaf litter contained 600-800 μg g^−1^ of cinnamic acids and comparable amounts of vanillin (Kuiters and Sarink, 1986). These concentrations would be expected to strongly inhibit growth of the bee parasites (IC50 195-414 μg mL^−1^ for cinnamic acid and 362-677 μg mL^−1^ for vanillin), but not *C. fasciculata* (extrapolated IC50 >1200 μg mL^−1^ for cinnamic acid and 1146-1487 μg mL^−1^ for vanillin). Prior studies documenting variation in tolerance of wood-derived tannins among mosquito species indicate that wood-associated phenolics in the larval habitat have exerted selection on mosquitoes (Rey *et al*., 2000). Our findings suggest that such compounds have shaped the mosquito-associated parasite community as well.

Although cinnamic acids also occur in honey and honey bee pollen (Mao *et al*., 2013)—where they likely result from degradation of cinnamic acid-spermidine conjugates that dominate the phytochemistry of plant pollen (Palmer-Young, Farrell, *et al*., 2019)—differences in digestive physiology between bees and mosquito are likely to result in lesser exposure of hindgut parasites to these compounds in bees. Mosquito larvae are filter feeders with a gut transit time of <1 h (Merritt *et al*., 1992), leaving little time for phytochemical metabolism in the midgut. In contrast, honey bees have a prolonged transit time (>12 h to reach the hindgut in newly emerged bees) (Gisder *et al*., 2020). As small-intestinal absorption of cinnamic acids is >95% in humans—where ingesta reach the large intestine in just 6-8 h (Olthof *et al*., 2001)—we hypothesize that minimal levels of these compounds reach the bee hindgut. As a result, *C. mellificae* and *L. passim* would have less direct exposure to these compounds than do *C. fasciculata* in mosquitoes.

### Comparisons with *Leishmania*

Both the two honey bee parasites and *C. fasciculata* showed high tolerance of some compounds previously shown to inhibit growth of *Leishmania*, which was particularly notable for flavonoids. This finding is consistent with the >30-fold higher tolerance to flavonoid-rich propolis extracts in *C. fasciculata* vs. *L. donovani* (Siheri 2016), which suggests ancestral tolerance to phytochemicals in the most recent common ancestor of these parasites that was selectively lost in *Leishmania* along with the acquisition of infectivity in mammals. One possible mechanistic explanation for this difference relates to the loss of catalase—an important enzyme for detoxification of reactive oxygen species (Kraeva *et al*., 2017; Horáková *et al*., 2020). A catalase of bacterial origin appears to have been present in the common ancestor of the Leishmaniinae subfamily before the divergence of the mammal-infecting *Leishmania* from the *Crithidia, Lotmaria*, and other insect-specific genera (Kraeva *et al*., 2017). This gene was likely under positive selection in the guts of nectar and especially pollen-feeding insects, where flavonoids can lead to reactive oxygen species-mediated stress that mediates their anti-leishmanial effects (Fonseca-Silva *et al*., 2011). However, based on the role of hydrogen peroxide in differentiation of *Leishmania* to the amastigote stage, reduced infectivity of catalase-expressing *Trypanosoma brucei*, and the absence of catalase in all mammal-infecting trypanosomatids, the presence of catalase appears to be incompatible with mammalian infection, which likely selected against this gene in *Leishmania* (Kraeva *et al*., 2017; Horáková *et al*., 2020). We hypothesize that the increased phytochemical sensitivity of *Leishmania* may reflect trade-offs that increase infectivity in mammals at the expense of tolerance to phytochemicals and associated reactive oxygen species in plant-feeding insects.

## CONCLUSIONS

The broad host range of insect-associated trypanosomatids makes their phytochemical tolerance and its ecological and evolutionary drivers a topic relevant for agriculture, conservation, and public health. Due to the high prevalence and effects of trypanosomatid infection in wild and managed honey, bumble, and solitary bees, development of phytochemical-based treatments or antiparasitic floral landscapes could contribute to sustainable pollination of crops and wildflowers. In addition, comparisons between parasite species that differ in phytochemical tolerance could illuminate the mechanisms that confer resistance to antiparasitic compounds, with potential to improve the efficacy of phytochemical-based treatments for neglected tropical diseases. Finally, trypanosomatids hitherto deemed insect-specific—including those with genetic identity to *C. fasiculata, C. mellificae*, and the close relative of bee trypanosomatids *Leptomonas seymouri* (Kraeva *et al*., 2015; Schwarz *et al*., 2015; Kaufer *et al*., 2017; Maruyama *et al*., 2019)—have been found in hematophagous trypanosome-vectoring tsetse flies (Votýpka *et al*., 2021), wild mammals (Dario *et al*., 2021), and humans suffering from visceral leishmaniasis-like illnesses (Maruyama *et al*., 2019). The infectivity of various insect-specific trypanosomatids in sand flies (Adler and Theodor, 1930) indicates that blood-seeking hosts may provide frequent opportunities for such taxa to ‘explore’ the mammalian niche (McGhee and Cosgrove, 1980; Lukeš *et al*., 2014). The zoonotic potential of these trypanosomatids portends the importance of their phytochemical tolerances for treatment of infection and environmentally friendly, landscape-based management of emerging parasites in vectors and reservoir hosts.

## ACKNOWLEDGMENTS

This project was supported by the USDA Agricultural Research Service Beltsville Bee Research Laboratory in house fund; a USDA-NIFA Pollinator Health Grant to JDE; and a North American Pollinator Protection Campaign Honey Bee Health Improvement Project Grant and an Eva Crane Trust Grant to ECPY and JDE. Funders had no role in study design, data collection and interpretation, or publication. We thank the ATCC, Ben Sadd, Quinn McFrederick, and Michael and Megan Povelones for sharing parasite strains and culturing advice; Steve Cook for use of spectrophotometers; Daniel Padfield for sharing R script; and anonymous reviewers for their service in improving the manuscript.

## DATA AVAILABILITY

All data are supplied in the Supplementary Information, Data S1.

## AUTHORS’ CONTRIBUTIONS

ECPY, JDE, and RSS conceived the study. ECPY conducted experiments, analyzed data, and drafted the manuscript with guidance from JDE, RSS, and YC. All authors revised the manuscript and gave approval for publication.

