## Supplementary information for "Punch in the gut: Parasite tolerance of phytochemicals reflects host diet"

Running title: Phytochemical niches of bee and mosquito parasites

Evan C Palmer-Young [1]*, Ryan S Schwarz [2], Jay D Evans [1]

[1] USDA-ARS Bee Research Lab, Beltsville, MD, USA

[2] Department of Biology, Fort Lewis College, Durango, CO, USA

### SUPPLEMENTARY FIGURES

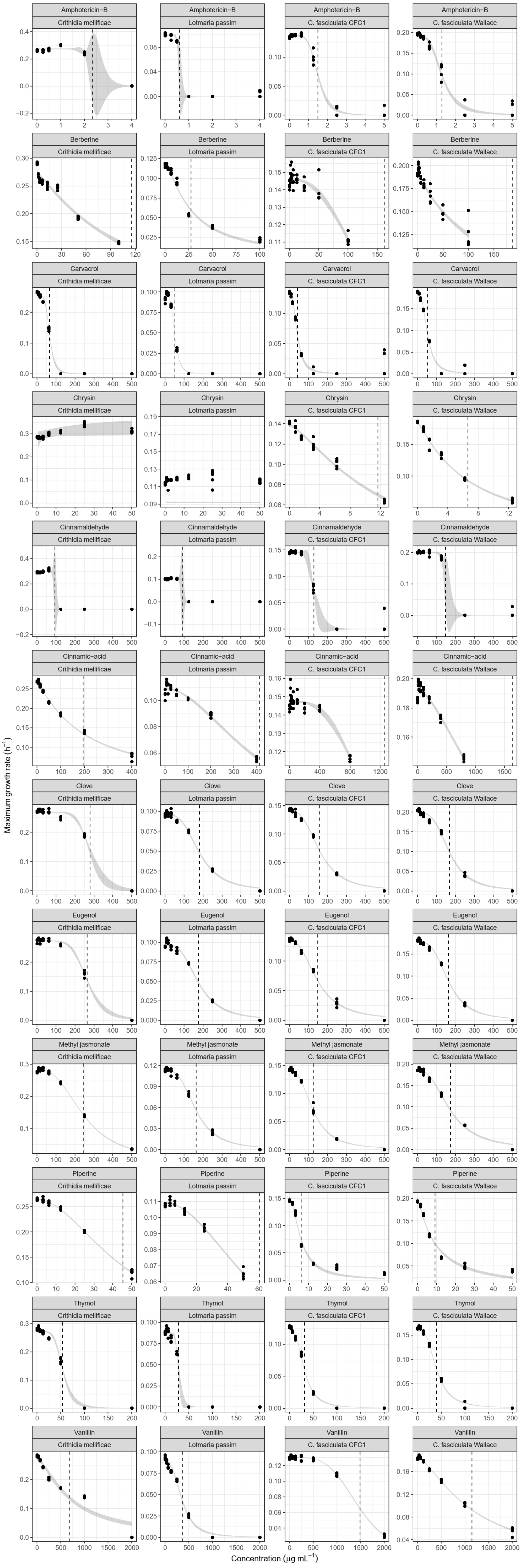
Amphotericin−B
Crithidia mellificae
Lotmaria passim
C. fasciculata CFC1
C. fasciculata Wallace
Berberine
Carvacrol
Chrysin
Cinnamaldehyde
Cinnamic−acid
Clove
Eugenol
Methyl jasmonate
Piperine
Thymol
Vanillin
Maximum growth rate (h$^{-1}$)
Concentration (μg mL$^{-1}$)

**Supplementary Figure 1. Dose-response curves for the 12 inhibitory compounds.** X-axis shows treatment concentration; Y-axis shows growth rate. Points indicate individual observations. Trendlines and shaded bands indicate model-predicted growth rates and standard errors. Vertical dashed lines show IC50 estimates, where applicable.

### SUPPLEMENTARY TABLES

**Supplementary Table 1.** Reported inhibitory concentrations (IC50) for the substances tested, plus the flavonoids apigenin and kaempferol and the esters chlorogenic and rosmarinic acids. Species are listed as in the original publications, but note that *L. chagasi* is considered synonymous with *L. infantum* (1). Although we attempted to test lemongrass extract, results were inconclusive due to poor solubility.

| Substance | Species | Stage | IC50 (µg/mL) | Assay | Duration (h) | Reference | Notes |
|---|---|---|---|---|---|---|---|
| Amphotericin-B | *Leishmania amazonensis* | P | 0.05 | Resazurin fluorescence | 24 | (2) | |
| Amphotericin-B | *Leishmania major* | P | 0.31 | Resazurin fluorescence | 72 | (3) | |
| Amphotericin-B | *Leishmania major* | P | 0.3 | Resazurin fluorescence | 72 | (4) | |
| Amphotericin-B | *Leishmania donovani* | P | 6.1 | MTT absorbance | 72 | (5) | |
| Amphotericin-B | *Leishmania infantum* | IA | 20.44 | ELISA | 48 | (6) | |
| Amphotericin-B | *Leishmania chagasi* | P | 0.51 | MTT absorbance | 72 | (7) | |
| Amphotericin-B | *Leishmania infantum* | P | 0.22 | MTT absorbance | 72 | (8) | |
| Amphotericin-B | *Leishmania major* | P | 0.8 | MTT absorbance | 72 | (8) | |
| Amphotericin-B | *Leishmania major* | P | 0.5 | Cell counts | 48 | (9) | |
| Amphotericin-B | *Leishmania major* | P | 0.96 | Cell counts | 48 | (9) | |
| Amphotericin-B | *Leishmania major* | IA | 0.2 | Proportion macrophages infected | 120 | (9) | |
| Amphotericin-B | *Leishmania major* | IA | 0.6 | Proportion macrophages infected | 120 | (9) | |
| Amphotericin-B | *Leishmania infantum* | IA | 20.44 | ELISA | 48 | (6) | |
| Berberine chloride | *Leishmania infantum* | P | 1.8 | MTS-PMS absorbance | 48 | (10) | |
| Berberine chloride | *Leishmania donovani* | IA | 2.5 | Cell counts | 72 | (11) | |
| Caffeic acid | *Leishmania amazonensis* | P | 0.9 | MTT absorbance | 72 | (12) | |
| Caffeic acid | *Leishmania amazonensis* | IA | 2.9 | Cell counts | 48 | (12) | |

| | | | | | | | |
|---|---|---|---|---|---|---|---|
| Carvacrol | *Leishmania amazonensis* | P | 25.4 | Resazurin fluorescence | 24 | (2) | |
| Carvacrol | *Leishmania infantum* | P | 7.35 | MTT absorbance | 72 | (8) | |
| Carvacrol | *Leishmania major* | P | 9.15 | MTT absorbance | 72 | (8) | |
| Carvacrol | *Leishmania chagasi* | P | 2.3 | MTT absorbance | 72 | (7) | |
| Carvacrol | *Leishmania chagasi* | P | 2.3 | MTT absorbance | 72 | (13) | |
| Carvacrol | *Leishmania amazonensis* | P | 15.3 | Nitrophenol phosphate hydrolysis | 72 | (14) | |
| Carvacrol | *Leishmania amazonensis* | P | 15.3 | MTT absorbance | 72 | (15) | |
| Carvacrol | *Leishmania amazonensis* | IA | 13.6 | Cell counts | 72 | (14) | |
| Carvacrol | *Leishmania infantum* | P | 9.8 | MTT absorbance | 24 | (16) | |
| Carvacrol | *Leishmania chagasi* | P | 27.3 | Cell counts | 72 | (17) | |
| Chrysin | *Leishmania donovani* | P | 13 | MTT absorbance | 24 | (18) | |
| Chrysin | *Leishmania mexicana* | P | 4.5 | Resazurin fluorescence | 72 | (19) | |
| Chrysin | *Leishmania donovani* | AA | 2.2 | Resazurin fluorescence | 72 | (20) | |
| Cinnam-aldehyde | *Leishmania mexicana* | P | 2.92 | Resazurin fluorescence | 72 | (21) | *Cinnamomum cassia* bark extract, 84% cinnamaldehyde |
| Cinnam-aldehyde | *Leishmania donovani* | P | 33.6 | Cell counts | 96 | (22) | *Cinnamomum cassia* bark dichloromethane extract, 84% cinnamaldehyde |
| Cinnamaldehyde | *Leishmania donovani* | IA | 14.06 | Cell counts | 48 | (22) | *Cinnamomum cassia* bark extract, 84% cinnamaldehyde |
| *trans*-cinnamic acid | *Leishmania amazonensis* | IA | 2.3 | Cell counts | 48 | (23) | |
| *trans*-cinnamic acid | *Leishmania mexicana* | P | 16.4 | Resazurin fluorescence | 72 | (19) | |
| Clove oil (*Eugenia* spp.) | *Leishmania donovani* | P | 21 | Cell counts | 96 | (24) | From *Szygium aromaticum* |
| Clove oil (*Eugenia* spp.) | *Leishmania donovani* | IA | 15.24 | Cell counts | 96 | (24) | From *Szygium aromaticum* |
| Clove oil (*Eugenia* spp.) | *Leishmania major* | P | 58.4 | Resazurin fluorescence | 72 | (4) | "Clove oil" |
| *p*-coumaric acid | *Leishmania amazonensis* | IA | 1.5 | Cell counts | 48 | (23) | |
| Eucalyptus oil (*Eucalyptus globulus*) | *Leishmania infantum* | P | 16.28 | MTT absorbance | 72 | (8) | |
| Eucalyptus oil (*Eucalyptus globulus*) | *Leishmania major* | P | 18.3 | MTT absorbance | 72 | (8) | |
| Eugenol | *Leishmania amazonensis* | P | 82.9 | Resazurin fluorescence | 24 | (2) | |
| Eugenol | *Leishmania amazonensis* | P | 80 | Cell viability | 1 | (25) | |
| Eugenol | *Leishmania infantum* | P | 56.13 | Bio-luminescence | 24 | (6) | |
| Eugenol | *Leishmania infantum* | IA | 20.81 | ELISA | 48 | (6) | |
| Eugenol | *Leishmania infantum chagasi* | P | 500 | Resazurin fluorescence | 24 | (26) | |
| Eugenol | *Leishmania infantum chagasi* | AA | 220 | Resazurin fluorescence | 24 | (26) | |
| Gallic acid | *Leishmania amazonensis* | IA | 2.6 | Cell counts | 48 | (23) | |

| | | | | | | | |
|---|---|---|---|---|---|---|---|
| Hesperetin | *Leishmania major* | P | 6.4 | Phospho-diesterase activity | 0.25 | (27) | |
| Hesperidin | *Leishmania donovani* | P | 622 | Cell counts | 24 | (28) | |
| Hesperidin | *Leishmania donovani* | IA | 171 | Cell counts | 24 | (28) | |
| Kaempferol | *Leishmania donovani* | AA | 2.9 | Resazurin fluorescence | 72 | (20) | |
| Kaempferol | *Leishmania donovani* | IA | 7.15 | Cell counts | 72 | (29) | |
| Kaempferol | *Leishmania peruviana* | P | 20.4 | Cell counts | 72 | (30) | |
| Kaempferol | *Leishmania braziliensis* | P | 15.3 | Cell counts | 72 | (30) | |
| Lemongrass oil (*Cymbopogon citratus*) | *Leishmania amazonensis* | P | 1.7 | Cell counts | 72 | (31) | |
| Lemongrass oil (*Cymbopogon citratus*) | *Leishmania amazonensis* | IA | 3.2 | Cell counts | 24 | (31) | |
| Linalool | *Leishmania amazonensis* | AA | 276.2 | Resazurin fluorescence | 24 | (32) | |
| Linalool | *Leishmania amazonensis* | P | 0.0043 | LD50 | 1 | (33) | |
| Linalool | *Leishmania amazonensis* | AA | 0.0155 | LD50 | 1 | (32) | |
| Linalool | *Leishmania infantum chagasi* | AA | 550 | Resazurin fluorescence | 24 | (26) | |
| Luteolin | *Leishmania donovani* | P | 3.6 | Cell counts | 24 | (34) | |
| Luteolin | *Leishmania donovani* | IA | 3.6 | Cell counts | 24 | (34) | |
| Luteolin | *Leishmania donovani* | AA | 0.7 | Resazurin fluorescence | 72 | (20) | |
| Methyl jasmonate | *Leishmania infantum* | IA | 24 | Cell counts | 72 | (35) | Methanolic extract of *Jasminum grandiflorum* |
| Methyl jasmonate | *Trypanosoma brucei brucei* | T | 31.2 | Cell counts | 72 | (35) | Methanolic extract of *Jasminum grandiflorum* |
| Methyl jasmonate | *Trypanosoma brucei rhodesiense* | T | 28.3 | Cell counts | 72 | (35) | Methanolic extract of *Jasminum grandiflorum* |
| Methyl jasmonate | *Trypanosoma cruzi* | IA | 29.3 | Galactosidase reporter activity | 168 | (35) | Methanolic extract of *Jasminum grandiflorum* |
| Methyl jasmonate | *Plasmodium falciparum* | | <112 | Intra-erythrocyte cell counts | 24 | (36) | |
| Methyl jasmonate | *Trichomonas vaginalis* | | 450 | Cell counts | 24 | (37) | |
| Methyl salicyclate | *Leishmania amazonensis* | P | 20.7 | MTT absorbance | 72 | (38) | Wintergreen (*Gualtheria fragrantissima*) essential oil (99.7% methyl salicylate) |

| | | | | | | | |
|---|---|---|---|---|---|---|---|
| Methyl salicyclate | *Leishmania amazonensis* | P | 32.2 | MTT absorbance | 72 | (38) | Birch bark (*Betula lenta*) essential oil (99.9% methyl salicylate) |
| Naringenin | *Leishmania donovani* | AA | 5 | Resazurin fluorescence | 72 | (20) | |
| Piperine | *Leishmania amazonensis* | P | 14.25 | Cell counts | 24 | (39) | |
| Quercetin | *Leishmania amazonensis* | P | 9.4 | Cell counts | 48 | (40) | |
| Quercetin | *Leishmania amazonensis* | P | 0.2 | MTT absorbance | 72 | (12) | |
| Quercetin | *Leishmania amazonensis* | IA | 1.3 | Cell counts | 48 | (12) | |
| Quercetin | *Leishmania amazonensis* | IA | 1 | Cell counts | 72 | (41) | |
| Quercetin | *Leishmania donovani* | P | 13.7 | Cell counts | 24 | (34) | |
| Quercetin | *Leishmania donovani* | AA | 1 | Resazurin fluorescence | 72 | (20) | |
| Quercetin | *Leishmania peruviana* | P | 16.1 | Cell counts | 72 | (30) | |
| Quercetin | *Leishmania braziliensis* | P | 20.7 | Cell counts | 72 | (30) | |
| Rutin | *Leishmania amazonensis* | IA | 2.7 | Cell counts | 48 | (23) | |
| Tea tree oil (*Melaleuca alternifolia*) | *Leishmania amazonensis* | P | 70.7 | MTT absorbance | 72 | (38) | |
| Tea tree oil (*Melaleuca alternifolia*) | *Leishmania major* | P | 403 | Resazurin fluorescence | 72 | (4) | |
| Thymol | *Leishmania amazonensis* | P | 26.8 | Resazurin fluorescence | 24 | (2) | |
| Thymol | *Leishmania infantum* | P | 12.85 | Bio-luminescence | 24 | (6) | |
| Thymol | *Leishmania infantum* | IA | 23.93 | ELISA | 48 | (6) | |
| Thymol | *Leishmania chagasi* | P | 9.8 | MTT absorbance | 72 | (7) | |
| Thymol | *Leishmania chagasi* | P | 9.8 | MTT absorbance | 72 | (13) | |
| Thymol | *Leishmania infantum* | P | 7.2 | MTT absorbance | 24 | (16) | |
| Thymol | *Leishmania amazonensis* | P | 19.5 | Cell counts | 48 | (42) | |
| Thymol | *Leishmania chagasi* | P | 65.2 | Cell counts | 72 | (17) | |
| Vanillin | *Leishmania donovani* | P | 0.5 | MTT absorbance | 72 | (43) | |
| Vanillin | *Leishmania donovani* | IA | 0.1 | Cell counts | 72 | (43) | |

Abbreviations:

P: Promastigote

AA: Axenic amastigote

IA: Intracellular amastigote

T: trypomastigote

MTS: [3-(4, 5 dimethyl-thiazol-2-yl) 5- (3-carboxymethoxyphenyl)-2-(4-sulphonyl)-2H-tetrazolium]"

PMS: phenazine methosulfate

MTT: 3-[4,5-dimethylthiazol-2-yl]-2,5-diphenyltetrazolium bromide

**Supplementary Table 2. Summary of 50% inhibitory concentrations** (IC50 (μg $mL^{-1}$) for *C. mellificae, L. passim,* and *C. fasciculata.* Columns "Lower" and "Upper" show 95% confidence intervals. "Conc.max" indicates maximum concentration tested. "Estimation" indicates whether the IC50 was within the tested concentration range ("Interpolated"), above the maximum concentration tested ("Extrapolated"), or not estimated due to insufficient inhibition.

| Class | Compound | Strain | IC50 | Lower | Upper | Conc.max | Estimation |
|---|---|---|---|---|---|---|---|
| Alkaloid | Berberine | Lotmaria passim | 27.43 | 25.06 | 29.8 | 100 | Interpolated |
| Alkaloid | Berberine | Crithidia mellificae | 115.81 | 101.12 | 130.51 | 100 | Extrapolated |
| Alkaloid | Berberine | C. fasciculata Wallace | 186.47 | 145.26 | 227.67 | 100 | Extrapolated |
| Alkaloid | Berberine | C. fasciculata CFC1 | >100 | | | 100 | Not estimated |
| Alkaloid | Piperine | Lotmaria passim | 60.56 | 57.98 | 63.14 | 50 | Extrapolated |
| Alkaloid | Piperine | Crithidia mellificae | 45.3 | 44.16 | 46.44 | 50 | Interpolated |
| Alkaloid | Piperine | C. fasciculata Wallace | 9.22 | 8.07 | 10.37 | 50 | Interpolated |
| Alkaloid | Piperine | C. fasciculata CFC1 | 6.09 | 5.5 | 6.68 | 50 | Interpolated |
| Benzenoid | Gallic acid | Crithidia mellificae | >500 | | | 500 | Not estimated |
| Benzenoid | Gallic acid | Lotmaria passim | >500 | | | 500 | Not estimated |
| Benzenoid | Gallic acid | C. fasciculata CFC1 | >500 | | | 500 | Not estimated |
| Benzenoid | Gallic acid | C. fasciculata Wallace | >500 | | | 500 | Not estimated |
| Benzenoid | Methyl salicylate | Crithidia mellificae | >500 | | | 500 | Not estimated |
| Benzenoid | Methyl salicylate | Lotmaria passim | >500 | | | 500 | Not estimated |
| Benzenoid | Methyl salicylate | C. fasciculata CFC1 | >500 | | | 500 | Not estimated |
| Benzenoid | Methyl salicylate | C. fasciculata Wallace | >500 | | | 500 | Not estimated |
| Benzenoid | Vanillin | Lotmaria passim | 362.3 | 339.38 | 385.23 | 2000 | Interpolated |
| Benzenoid | Vanillin | Crithidia mellificae | 677.08 | 545.03 | 809.14 | 2000 | Interpolated |
| Benzenoid | Vanillin | C. fasciculata Wallace | 1146.06 | 1105.02 | 1187.1 | 2000 | Interpolated |
| Benzenoid | Vanillin | C. fasciculata CFC1 | 1487.36 | 1463.1 | 1511.61 | 2000 | Interpolated |
| Extract | Clove | Lotmaria passim | 181.09 | 173.77 | 188.4 | 500 | Interpolated |
| Extract | Clove | Crithidia mellificae | 279.83 | 266 | 293.65 | 500 | Interpolated |
| Extract | Clove | C. fasciculata Wallace | 168.29 | 161.04 | 175.54 | 500 | Interpolated |
| Extract | Clove | C. fasciculata CFC1 | 159.64 | 154.39 | 164.89 | 500 | Interpolated |
| Extract | Eucalyptus | Crithidia mellificae | >500 | | | 500 | Not estimated |
| Extract | Eucalyptus | Lotmaria passim | >500 | | | 500 | Not estimated |
| Extract | Eucalyptus | C. fasciculata CFC1 | >500 | | | 500 | Not estimated |
| Extract | Eucalyptus | C. fasciculata Wallace | >500 | | | 500 | Not estimated |
| Extract | Tea tree | Crithidia mellificae | >500 | | | 500 | Not estimated |
| Extract | Tea tree | Lotmaria passim | >500 | | | 500 | Not estimated |
| Extract | Tea tree | C. fasciculata CFC1 | >500 | | | 500 | Not estimated |
| Extract | Tea tree | C. fasciculata Wallace | >500 | | | 500 | Not estimated |
| Flavonoid | Chrysin | Lotmaria passim | >50 | | | 50 | Not estimated |
| Flavonoid | Chrysin | Crithidia mellificae | >50 | | | 50 | Not estimated |
| Flavonoid | Chrysin | C. fasciculata Wallace | 6.65 | 6.18 | 7.13 | 12.5 | Interpolated |
| Flavonoid | Chrysin | C. fasciculata CFC1 | 11.69 | 10.83 | 12.56 | 12.5 | Interpolated |
| Flavonoid | Hesperetin | Crithidia mellificae | >50 | | | 50 | Not estimated |
| Flavonoid | Hesperetin | Lotmaria passim | >50 | | | 50 | Not estimated |

| | | | | | | | |
|---|---|---|---|---|---|---|---|
| Flavonoid | Hesperetin | C. fasciculata CFC1 | >50 | | | 50 | Not estimated |
| Flavonoid | Hesperetin | C. fasciculata Wallace | >50 | | | 50 | Not estimated |
| Flavonoid | Luteolin | Crithidia mellificae | >50 | | | 50 | Not estimated |
| Flavonoid | Luteolin | Lotmaria passim | >50 | | | 50 | Not estimated |
| Flavonoid | Luteolin | C. fasciculata CFC1 | >50 | | | 50 | Not estimated |
| Flavonoid | Luteolin | C. fasciculata Wallace | >50 | | | 50 | Not estimated |
| Flavonoid | Naringenin | Crithidia mellificae | >50 | | | 50 | Not estimated |
| Flavonoid | Naringenin | Lotmaria passim | >50 | | | 50 | Not estimated |
| Flavonoid | Naringenin | C. fasciculata CFC1 | >50 | | | 50 | Not estimated |
| Flavonoid | Naringenin | C. fasciculata Wallace | >50 | | | 50 | Not estimated |
| Flavonoid | Quercetin | Crithidia mellificae | >50 | | | 50 | Not estimated |
| Flavonoid | Quercetin | Lotmaria passim | >50 | | | 50 | Not estimated |
| Flavonoid | Quercetin | C. fasciculata CFC1 | >50 | | | 50 | Not estimated |
| Flavonoid | Quercetin | C. fasciculata Wallace | >50 | | | 50 | Not estimated |
| Flavonoid glycoside | Hesperidin | Crithidia mellificae | >50 | | | 50 | Not estimated |
| Flavonoid glycoside | Hesperidin | Lotmaria passim | >50 | | | 50 | Not estimated |
| Flavonoid glycoside | Hesperidin | C. fasciculata CFC1 | >50 | | | 50 | Not estimated |
| Flavonoid glycoside | Hesperidin | C. fasciculata Wallace | >50 | | | 50 | Not estimated |
| Flavonoid glycoside | Rutin | Crithidia mellificae | >200 | | | 200 | Not estimated |
| Flavonoid glycoside | Rutin | Lotmaria passim | >200 | | | 200 | Not estimated |
| Flavonoid glycoside | Rutin | C. fasciculata CFC1 | >200 | | | 200 | Not estimated |
| Flavonoid glycoside | Rutin | C. fasciculata Wallace | >200 | | | 200 | Not estimated |
| Oxylipin | Methyl jasmonate | Lotmaria passim | 162.54 | 157.28 | 167.79 | 500 | Interpolated |
| Oxylipin | Methyl jasmonate | Crithidia mellificae | 245.74 | 241.27 | 250.22 | 500 | Interpolated |
| Oxylipin | Methyl jasmonate | C. fasciculata Wallace | 172.3 | 164.28 | 180.32 | 500 | Interpolated |
| Oxylipin | Methyl jasmonate | C. fasciculata CFC1 | 124.02 | 120.18 | 127.86 | 500 | Interpolated |
| Phenylpropanoid | Caffeic acid | Crithidia mellificae | >400 | | | 400 | Not estimated |
| Phenylpropanoid | Caffeic acid | Lotmaria passim | >400 | | | 400 | Not estimated |
| Phenylpropanoid | Caffeic acid | C. fasciculata CFC1 | >400 | | | 400 | Not estimated |
| Phenylpropanoid | Caffeic acid | C. fasciculata Wallace | >400 | | | 400 | Not estimated |
| Phenylpropanoid | Cinnamaldehyde | Lotmaria passim | 90.27 | 28.75 | 151.79 | 500 | Interpolated |
| Phenylpropanoid | Cinnamaldehyde | Crithidia mellificae | 94.78 | 59.7 | 129.86 | 500 | Interpolated |
| Phenylpropanoid | Cinnamaldehyde | C. fasciculata Wallace | 148.89 | 106.05 | 191.72 | 500 | Interpolated |
| Phenylpropanoid | Cinnamaldehyde | C. fasciculata CFC1 | 127.73 | 123.35 | 132.12 | 500 | Interpolated |
| Phenylpropanoid | Cinnamic acid | Lotmaria passim | 413.84 | 389.54 | 438.15 | 400 | Extrapolated |
| Phenylpropanoid | Cinnamic acid | Crithidia mellificae | 194.89 | 184.35 | 205.44 | 400 | Interpolated |
| Phenylpropanoid | Cinnamic acid | C. fasciculata Wallace | >800 | | | 800 | Not estimated |
| Phenylpropanoid | Cinnamic acid | C. fasciculata CFC1 | >800 | | | 800 | Not estimated |
| Phenylpropanoid | Coumaric acid | Crithidia mellificae | >400 | | | 400 | Not estimated |
| Phenylpropanoid | Coumaric acid | Lotmaria passim | >400 | | | 400 | Not estimated |
| Phenylpropanoid | Coumaric acid | C. fasciculata CFC1 | >400 | | | 400 | Not estimated |
| Phenylpropanoid | Coumaric acid | C. fasciculata Wallace | >400 | | | 400 | Not estimated |
| Phenylpropanoid | Eugenol | Lotmaria passim | 175.95 | 168.35 | 183.54 | 500 | Interpolated |
| Phenylpropanoid | Eugenol | Crithidia mellificae | 264.23 | 258.86 | 269.6 | 500 | Interpolated |

| | | | | | | | |
|---|---|---|---|---|---|---|---|
| Phenylpropanoid | Eugenol | C. fasciculata Wallace | 163.53 | 158.32 | 168.74 | 500 | Interpolated |
| Phenylpropanoid | Eugenol | C. fasciculata CFC1 | 146.89 | 140.87 | 152.91 | 500 | Interpolated |
| Polyene | Amphotericin B | Crithidia mellificae | 2.32 | 1.07 | 3.58 | 4 | Interpolated |
| Polyene | Amphotericin B | Lotmaria passim | 0.59 | 0.41 | 0.77 | 4 | Interpolated |
| Polyene | Amphotericin B | C. fasciculata Wallace | 1.29 | 1.19 | 1.38 | 5 | Interpolated |
| Polyene | Amphotericin B | C. fasciculata CFC1 | 1.5 | 1.44 | 1.56 | 5 | Interpolated |
| Terpenoid | Carvacrol | Lotmaria passim | 50.61 | 48.96 | 52.27 | 500 | Interpolated |
| Terpenoid | Carvacrol | Crithidia mellificae | 65.7 | 64.05 | 67.35 | 500 | Interpolated |
| Terpenoid | Carvacrol | C. fasciculata Wallace | 52.36 | 48.98 | 55.74 | 500 | Interpolated |
| Terpenoid | Carvacrol | C. fasciculata CFC1 | 41.08 | 37.21 | 44.95 | 500 | Interpolated |
| Terpenoid | Linalool | Crithidia mellificae | >500 | | | 500 | Not estimated |
| Terpenoid | Linalool | Lotmaria passim | >500 | | | 500 | Not estimated |
| Terpenoid | Linalool | C. fasciculata CFC1 | >500 | | | 500 | Not estimated |
| Terpenoid | Linalool | C. fasciculata Wallace | >500 | | | 500 | Not estimated |
| Terpenoid | Thymol | Lotmaria passim | 28.3 | 25.84 | 30.76 | 200 | Interpolated |
| Terpenoid | Thymol | Crithidia mellificae | 54.1 | 52.39 | 55.8 | 200 | Interpolated |
| Terpenoid | Thymol | C. fasciculata Wallace | 39.56 | 38.13 | 40.98 | 200 | Interpolated |
| Terpenoid | Thymol | C. fasciculata CFC1 | 31.01 | 29.76 | 32.25 | 200 | Interpolated |

### SUPPLEMENTARY DATA

**Zipped folder with data spreadsheets** for *Leishmania* inhibitory concentrations (**leishmania_ic50**) and growth rates (**rates.selected**) and IC50 estimates (**df.ic50)** for the parasites tested here.
